# Glucocorticoids inhibit macrophage differentiation towards a pro-inflammatory phenotype upon wounding without affecting their migration

**DOI:** 10.1101/473926

**Authors:** Yufei Xie, Sofie Tolmeijer, Jelle Oskam, Tijs Tonkens, Annemarie H. Meijer, Marcel J.M Schaaf

**Affiliations:** Institute of Biology, Leiden University, Leiden, The Netherlands

**Keywords:** glucocorticoids, inflammation, macrophage differentiation, leukocyte migration, zebrafish, tail amputation

## Abstract

Glucocorticoid drugs are widely used to treat immune-related diseases, but their use is limited by side effects and by resistance, which especially occurs in macrophage-dominated diseases. In order to improve glucocorticoid therapies, more research is required into the mechanisms of glucocorticoid action. In the present study, we have used a zebrafish model for inflammation to study glucocorticoid effects on the innate immune response. In zebrafish larvae, the migration of neutrophils towards a site of injury is inhibited by the synthetic glucocorticoid beclomethasone, while migration of macrophages is glucocorticoid resistant. We show that wounding-induced increases in expression of genes encoding neutrophil-specific chemoattractants (Il8 and Cxcl18b) are attenuated by beclomethasone, but that beclomethasone does not attenuate the induction of the genes encoding Ccl2 and Cxcl11aa, which we show to be required for macrophage recruitment. RNA sequencing on Fluorescence-Activated Cell Sorting (FACS)-sorted macrophages showed that the vast majority of the wounding-induced transcriptional changes in these cells are inhibited by beclomethasone, whereas a small subset is glucocorticoid-insensitive. As a result, beclomethasone decreases the number of macrophages that differentiate towards a pro-inflammatory (M1) phenotype, which we demonstrated using a *tnfa:eGFP-F* reporter line and analysis of macrophage morphology. We conclude that the glucocorticoid resistance of the wounding-induced macrophage migration is due to the insensitivity of the induction of macrophage-specific chemoattractants to glucocorticoid inhibition, which may explain the relative resistance of macrophage-dominated diseases to glucocorticoid therapy. However, the induction of pro-inflammatory genes in macrophages is strongly attenuated, which inhibits their differentiation to an M1 phenotype.

**Summary statement:** In a zebrafish model for inflammation, glucocorticoids do not affect the migration of macrophages, but inhibit differentiation towards an M1 phenotype, by strongly attenuating transcriptional responses in these cells.

## Introduction

Glucocorticoids (GCs) are a class of steroid hormones secreted by the adrenal gland, and the main endogenous glucocorticoid in our body is cortisol (Chrousos 1995; Oakley and Cidlowski, 2013; Tsigos and Chrousos, 2002). Glucocorticoids regulate a wide variety of systems in our body, including the immune, metabolic, reproductive, cardiovascular and central nervous system (Chrousos and Kino, 2005; Heitzer et al., 2007; Ramamoorthy and Cidlowski, 2013; Revollo and Cidlowski, 2009). Due to their potent and well-established immunosuppressive effects, they are often prescribed to treat various immune-related diseases, including asthma, rheumatoid arthritis, dermatitis, leukemia, and several autoimmune diseases (Barnes, 2011; Busillo and Cidlowski, 2013). However, their clinical use is limited by two issues. First, chronic glucocorticoid therapy can lead to severe side effects, like osteoporosis, muscle weakness, diabetes, infection, and hypertension (Moghadam-Kia and Werth, 2010). Second, resistance to glucocorticoid drug treatment occurs in a large number (∼10-30%) of patients (Barnes and Adcock, 2009; Barnes et al., 2004). In order to develop novel glucocorticoid therapies that overcome these barriers and retain their therapeutic efficacy, more insight into the molecular and cellular mechanisms of glucocorticoid modulation of the immune response is required.

Glucocorticoids exert their function through an intracellular receptor, the glucocorticoid receptor (GR) (Bamberger et al., 1996), which acts as a transcription factor, altering the transcription of a plethora of genes. The GR modulates the transcription of genes by several mechanisms (Ratman et al., 2013). It can bind directly to DNA, to glucocorticoid response elements (GRE) and enhance transcription upon recruitment of transcriptional cofactors. In contrast, binding to negative GREs (nGREs) has been shown to repress gene transcription (Surjit et al., 2011). Alternatively, the GR can bind indirectly to DNA through interaction with other transcription factors, like AP-1, NF-κB or STAT3. Through this ‘tethering’, it modulates the activity of these factors.

The tethering mechanism of the GR, resulting in the inhibition of transcription of immune-activating genes, is generally considered to be the main mechanism by which glucocorticoids exert their anti-inflammatory actions (Reichardt et al., 2001). For example, TNF- or LPS-induced transcriptional responses in cultured cells can be repressed through tethering of the NF-κB subunit p65 (Kuznetsova et al., 2015; Ogawa et al., 2005; Rao et al., 2011; Sacta et al., 2018). Other mechanisms, like the activation of anti-inflammatory genes through GRE binding, and a reduction of NF-κB recruitment, contribute to the anti-inflammatory actions of GR as well, but the exact role of these mechanisms has not been fully established (Hubner et al., 2015; Oh et al., 2017). Through these mechanisms, glucocorticoids exert strong suppressive effects on the inflammatory response (Smoak and Cidlowski, 2004). At the initial stage of this response, they dampen signaling pathways downstream from Toll-like receptors (TLRs), inhibit the induction of genes encoding cytokines, upregulate the expression of anti-inflammatory proteins, and inhibit the generation of prostaglandins and leukotrienes (Busillo and Cidlowski, 2013; Coutinho and Chapman, 2011). In addition, they reduce the blood flow to the inflamed tissue and inhibit vascular leakage. At subsequent stages, glucocorticoids attenuate the production of chemokines and adhesion molecules, thereby reducing leukocyte extravasation and migration towards the inflamed site (Coutinho and Chapman, 2011; Smoak and Cidlowski, 2004).

It has become clear that glucocorticoid action on the immune system is highly complex and requires further investigation. A complicating factor is that the effects of glucocorticoids have been shown to be highly cell type-specific. Whereas they induce apoptosis of eosinophils and basophils, they promote the survival and proliferation of neutrophils (Meagher et al., 1996; Yoshimura et al., 2001). In monocytes, they induce an anti-inflammatory phenotype with increased mobility and phagocytic capacity (Ehrchen et al., 2007). Macrophages can be divided into two functional phenotypes (Nguyen-Chi et al., 2015), a pro-inflammatory (M1) phenotype which contributes to the inflammatory response (Baschant and Tuckermann, 2010), and an anti-inflammatory (M2) phenotype which displays enhanced phagocytic activity and is mostly involved in the resolution of inflammation (Busillo and Cidlowski, 2013). Glucocorticoids have been shown to affect the differentiation of macrophages towards these phenotypes (Hofkens et al., 2013; Tu et al., 2017). In addition to the cell type-specificity of glucocorticoid actions, it has become clear that the transcriptional regulation of immune-activating genes by the GR is not strictly suppressive (Cruz-Topete and Cidlowski, 2015).

Upregulation of various pro-inflammatory genes after glucocorticoid treatment has been observed in several cell types (Busillo et al., 2011; Chinenov and Rogatsky, 2007; Ding et al., 2010; Galon et al., 2002; Lannan et al., 2012), and GR has been shown to activate pro-inflammatory genes in synergy with other signaling pathways (Anna et al., 2012; Langlais et al., 2008; Langlais et al., 2012). In addition, some genes that are induced upon TNF or LPS treatment appear to be insensitive to the repressive action of GR (Kuznetsova et al., 2015; Ogawa et al., 2005; Rao et al., 2011; Sacta et al., 2018).

In the present study, we have used the zebrafish as an *in vivo* model to study glucocorticoid effects on the inflammatory response. The immune system of the zebrafish is highly similar to that of humans. Like humans, the zebrafish has a thymus, innate immune cells (macrophages, neutrophils) and adaptive immune cells (T cells and B cells), and cells that bridge innate and adaptive immunity (dendritic cells) (Lewis et al., 2014; Masud et al., 2017; Sullivan et al., 2017). Besides, the innate immune system of zebrafish develops within a few days after fertilization, while the adaptive immune system only matures after two weeks, which means the innate immune system can be studied separately in larvae (Masud et al., 2017; Trede et al., 2004). Zebrafish larvae are widely used as a model system to study the inflammatory response (Enyedi et al., 2016; Oehlers et al., 2017; Powell et al., 2017). Tail wounding-induced inflammation in zebrafish larvae is a well-established model in which amputation of the tail triggers the expression of many pro-inflammatory molecules and the recruitment of innate immune cells (neutrophils and macrophages) towards the wounded area (Renshaw et al., 2006; Roehl, 2018). This model enables the investigation of cell-type specific inflammatory responses *in vivo* and has been widely used for research on leukocyte migration and infiltration and anti-inflammatory drug screening (Niethammer et al., 2009; Robertson et al., 2016; Yoo et al., 2011).

The zebrafish Gr is highly similar to its human equivalent in structure and function (Chatzopoulou et al., 2015; Schaaf et al., 2008; Stolte et al., 2006). This makes the zebrafish a valuable model to study the molecular mechanisms of glucocorticoid action *in vivo* (Alsop and Vijayan, 2008; Schaaf et al., 2008; Schaaf et al., 2009). In previous work, we have studied the anti-inflammatory effects of glucocorticoids using the tail amputation model and found that glucocorticoid treatment attenuates the vast majority amputation-induced changes in gene expression (Chatzopoulou et al., 2016). In addition, we observed that the recruitment of neutrophils to the wounded area is inhibited by glucocorticoids, but that the migration of macrophages is resistant to glucocorticoid treatment (Chatzopoulou et al., 2016; Mathew et al., 2007; Zhang et al., 2008).

It has been shown that glucocorticoids are less effective in the treatment of inflammatory diseases dominated by macrophages, like chronic obstructive pulmonary disease (COPD), but the mechanisms underlying the limited responsiveness to glucocorticoid treatment remain poorly understood (Hakim et al., 2012). Therefore, in the present study, we sought to find a mechanistic explanation for our finding that glucocorticoids do not inhibit amputation-induced macrophage migration. We demonstrate that the induction of genes encoding chemo-attractants involved in macrophage recruitment is insensitive to glucocorticoid treatment, providing an explanation for the insensitivity of macrophage migration to glucocorticoids. However, macrophages should not be considered a generally glucocorticoid-insensitive cell type, since in these cells, glucocorticoids attenuated almost all amputation-induced changes in gene expression, which involved a wide variety of immunity- and metabolism-related genes. Through this modulation of the transcriptional response, glucocorticoids inhibit the differentiation of macrophages to a pro-inflammatory (M1) phenotype.

## Results

### Beclomethasone inhibits migration of neutrophils, but leaves macrophage migration unaffected

Using tail amputation in 3 dpf zebrafish larvae as a model for inflammation, we studied the effect of the synthetic glucocorticoid beclomethasone on the migration of leukocytes towards a site of injury. To quantitate the migration of neutrophils and macrophages, we counted the number of these innate immune cells in a defined area of the tail between 1.5 and 12 hours post amputation (hpa) (Figure 1A). The results of this quantitation showed that for the control group the average number of macrophages present in the wounded area increased from 37.0±3.5 to 48.7±4.1 cells between 1.5 hpa and 12 hpa (Figure 1B). No significant effect of beclomethasone on macrophage migration was observed (from 37.6±2.8 to 41.4±2.5 for the beclomethasone-treated group), in line with our previous results (Chatzopoulou et al., 2016). For neutrophils, in the control group, a number of 17.7±2.0 was observed at 1.5 hpa, and their number reached a peak of 35.1±4.1 at around 5 hpa, then decreased and reached a level of 32.7±3.3 at 9 hpa which remained relatively constant until 12 hpa (Figure 1C). In the beclomethasone-treated group, a lower number of recruited neutrophils was observed in the wounded area at 5 hpa (22.8±1.9).

**Figure 1.**
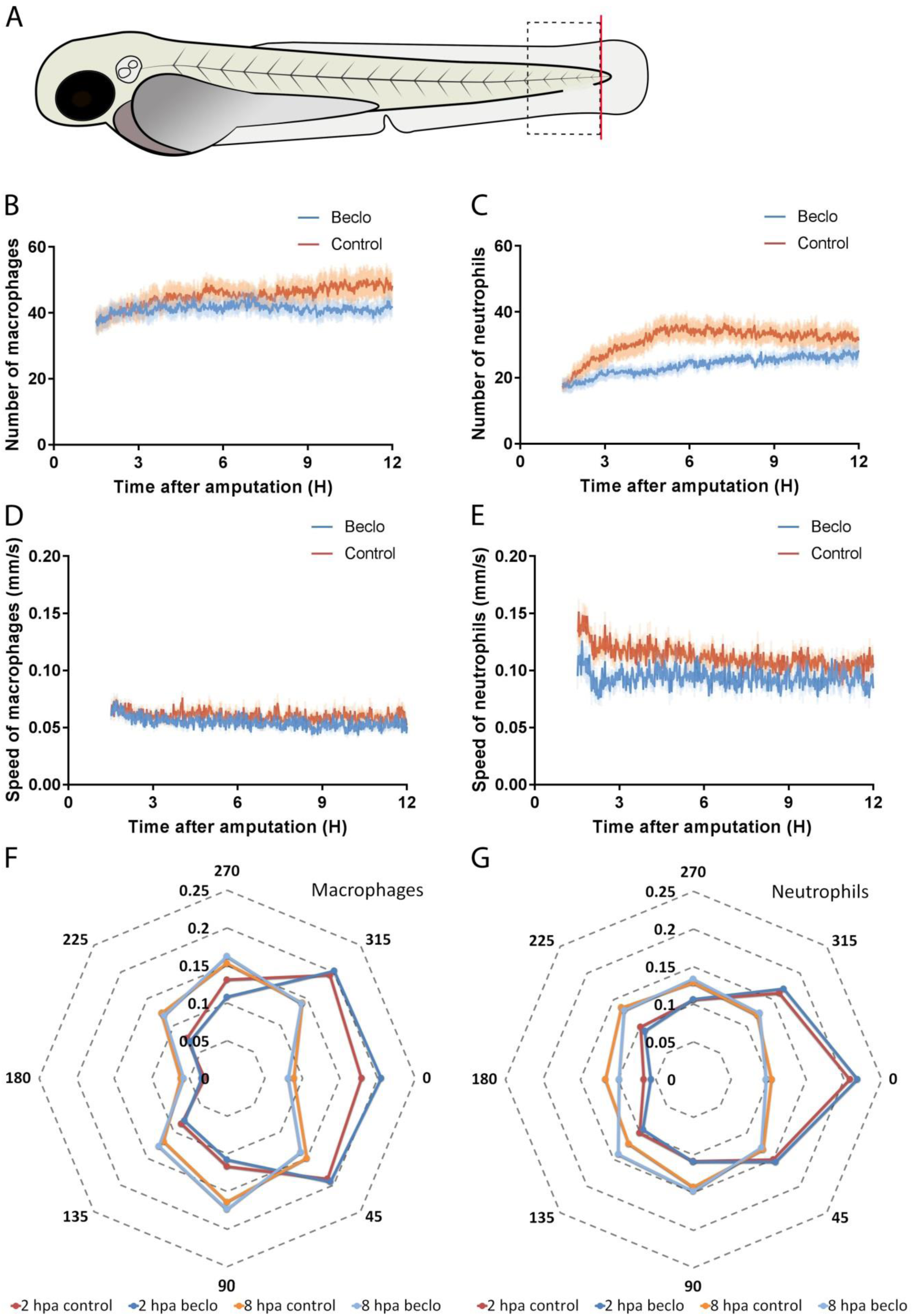
Live imaging and tracking of migrating macrophages and neutrophils. In the tail amputation assay. A. Schematic drawing of a zebrafish larva at 3 dpf. The red line shows the site of amputation. The black dashed box shows the area in which cells were counted to quantitate the recruitment. B-C. The number of macrophages (B) and neutrophils (C) recruited to the wounded area from 1.5 hpa to 12 hpa in 3 dpf larvae. No significant difference was observed for the number of recruited macrophages. A significantly reduced number of neutrophils were recruited in the beclomethasone treated group compared to the control group at 5 hpa. D-E. The velocity of macrophages (D) and neutrophils (E). No significant difference was observed for the velocity of macrophages. At 2 and 3 hpa, the velocity of neutrophils in the beclomethasone-treated group was significantly lower than the velocity in the control group. F-G. The directionality of recruited macrophages (F) and neutrophils (G) at 2 hpa and 8 hpa. The circular x axis indicates the different angles made by cells, classified into 8 categories. Category 0 including angles between 22.5 to −22.5 degrees and represents the direction towards the wound. The y axis indicates the size of the fraction of cells occurring within a category in that hour. No difference was observed between the control and beclomethasone treated groups.

To analyze the effects of beclomethasone in more detail, we quantified the velocity and directionality of migrating macrophages and neutrophils. The data showed that during the entire time lapse, the velocity of the macrophages fluctuated around 0.06 mm/s for both the control and the beclomethasone-treated group (Figure 1D). For neutrophils, the velocity peaked at 1.5 hpa (0.135±0.009 mm/s for the control group and 0.095±0.012 mm/s for the beclomethasone-treated group) and decreased slowly afterwards (Figure 1E). At 2 hpa and 3 hpa, the velocity of neutrophils in the beclomethasone-treated group was significantly lower compared to the control group.

In addition, we measured the direction in which the macrophages and neutrophils moved and plotted the distribution of these directions measured at 2 and 8 hours after amputation (Figures 1F-G). The results showed that at none of these time points beclomethasone affected the directionality of either macrophages or neutrophils. At 2 hpa, most of the macrophages (∼60%) moved towards the wounded area (angles 292.5°- 360°, and 0°- 67.5°) (Figure 1F). Only less than 20% of them moved in the opposite direction (angles 112.5°- 247.5°). At 8 hpa, the percentage of macrophages that moved towards the wounded area in the control and beclomethasone-treated group decreased to ∼40%. For the neutrophils, the directionality showed a similar trend (Figure 1G). At 2 hpa, over 50% of the neutrophils moved towards the wounded area in both the control group and the beclomethasone-treated group, while at 8 hpa, this percentage decreased to ∼35%. In conclusion, beclomethasone does not affect any of the migration parameters of macrophages but reduces the number of recruited neutrophils and their velocity.

### Beclomethasone inhibits the induction of chemoattractants for macrophages

To unravel the molecular mechanisms underlying the difference between the effect of beclomethasone on macrophage and neutrophil migration, we first studied the expression of chemo-attractants that are known to be involved in the migration of these leukocytes. According to previous studies on leukocyte migration and infiltration, Ccl2 and Cxcl11aa are two of the key chemokines that stimulate the migration of macrophages, while Il8 and Cxcl18b are important for the stimulation of neutrophil migration (Cambier et al., 2017; de Oliveira et al., 2013; de Oliveira et al., 2016; Deshmane et al., 2009; Huber et al., 1991; Torraca et al., 2015; Torraca et al., 2017). Using qPCR on RNA samples from whole larvae, we determined the expression levels of the genes encoding these four chemo-attractants (*ccl2*, *cxcl11aa, il8* and *cxcl18b)* at different time points after amputation, (Figure 2A-D). The results showed that at 4 hpa, the mRNA level of all four chemo-attractants was increased by amputation. At 2 hpa, the expression of *ccl2*, *cxcl11aa*, and *cxcl18b* was increased, and at 8 hpa the expression of *ccl2*, *il8* and *cxcl18b* showed an increase. In the presence of beclomethasone, amputation induced a smaller increase in *il8* and *cxcl18b* expression but the increase in the expression of *ccl2* and *cxcl11aa* was not inhibited.

**Figure 2.**
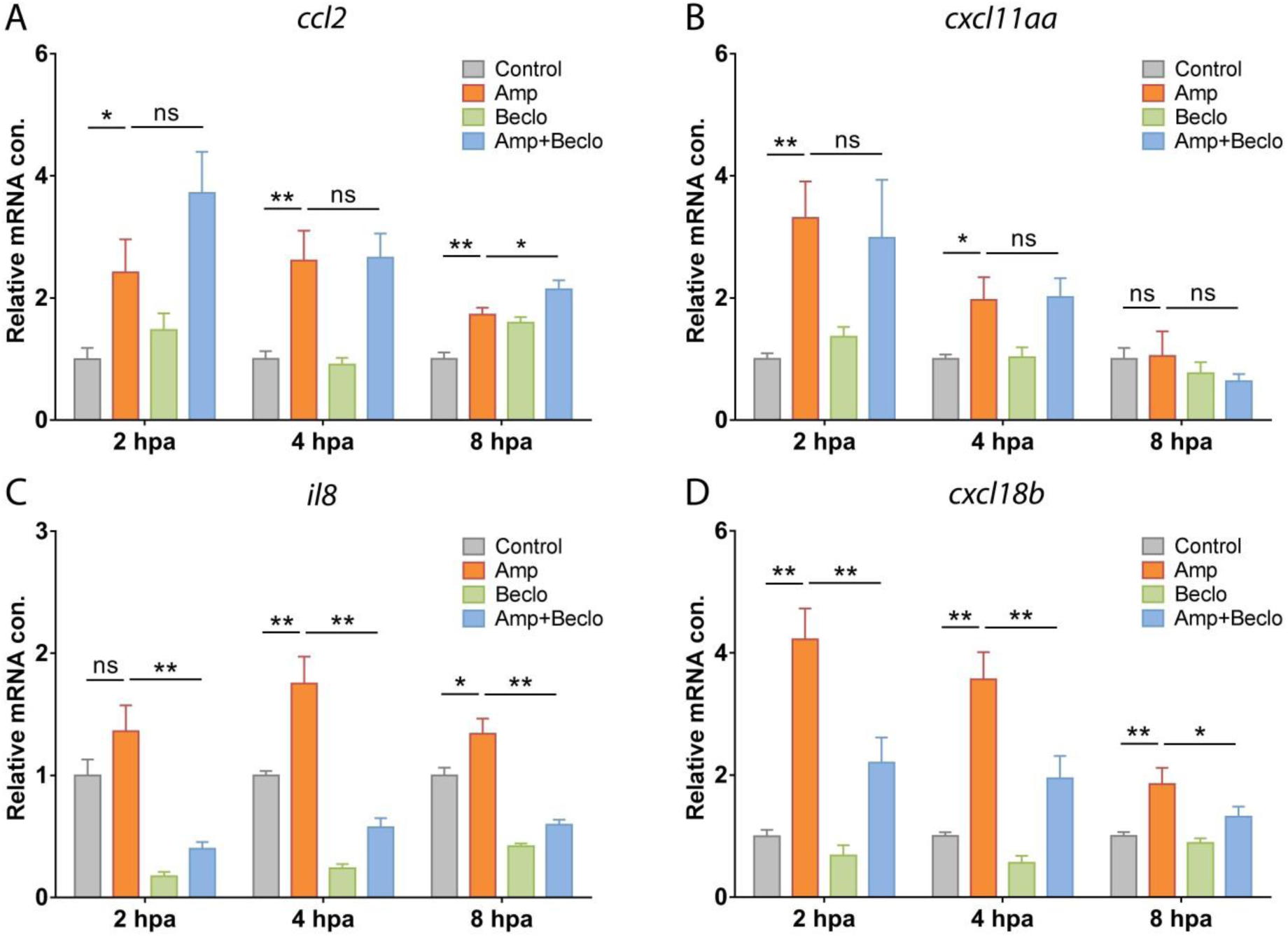
Expression levels of genes encoding chemo-attractants Ccl2 (A), Cxcl11aa (B), Il8 (C), and Cxcl18b (D) in whole larvae at 2hpa, 4hpa and 8hpa, determined by qPCR. Statistical analysis showed that *ccl2* and *cxcl11aa* mRNA levels were significantly increased by amputation (Amp) and that the combined amputation/beclomethasone (Amp+Beclo) treatment resulted in a similar level of regulation. Expression levels of *il8*, and *cxcl18b* showed a significant increase upon amputation, and this effect was lower upon the combined treatment. Values shown are the means ± s.e.m. of three independent experiments. Statistical significance is indicated by: ns, non-significant; * P<0.05; ** P<0.01.

To demonstrate that the chemo-attractants Ccl2 and Cxcl11aa are required for macrophage recruitment in this tail amputation model, we analyzed their role in macrophage migration in our model. We used a previously described morpholino to create a knockdown of Ccr2, the receptor of Ccl2, in zebrafish larvae(Cambier et al., 2017; Cambier et al., 2014). In the *ccr2* morphants, a significantly decreased number of recruited macrophages was observed in the wounded area at 4 hpa (Figure 3A, C, D). However, the number of recruited neutrophils was identical to the number in the wild type (Figure 3B, E, F). For the receptor of Cxcl11aa, Cxcr3.2, we used a mutant fish line (Torraca et al., 2015). The *cxcr3.*2 −/− larvae showed significantly decreased numbers of both macrophages (Figure 4A, C, D) and neutrophils (Figure 4B, E, F) recruited to the wounded area.

**Figure 3.**
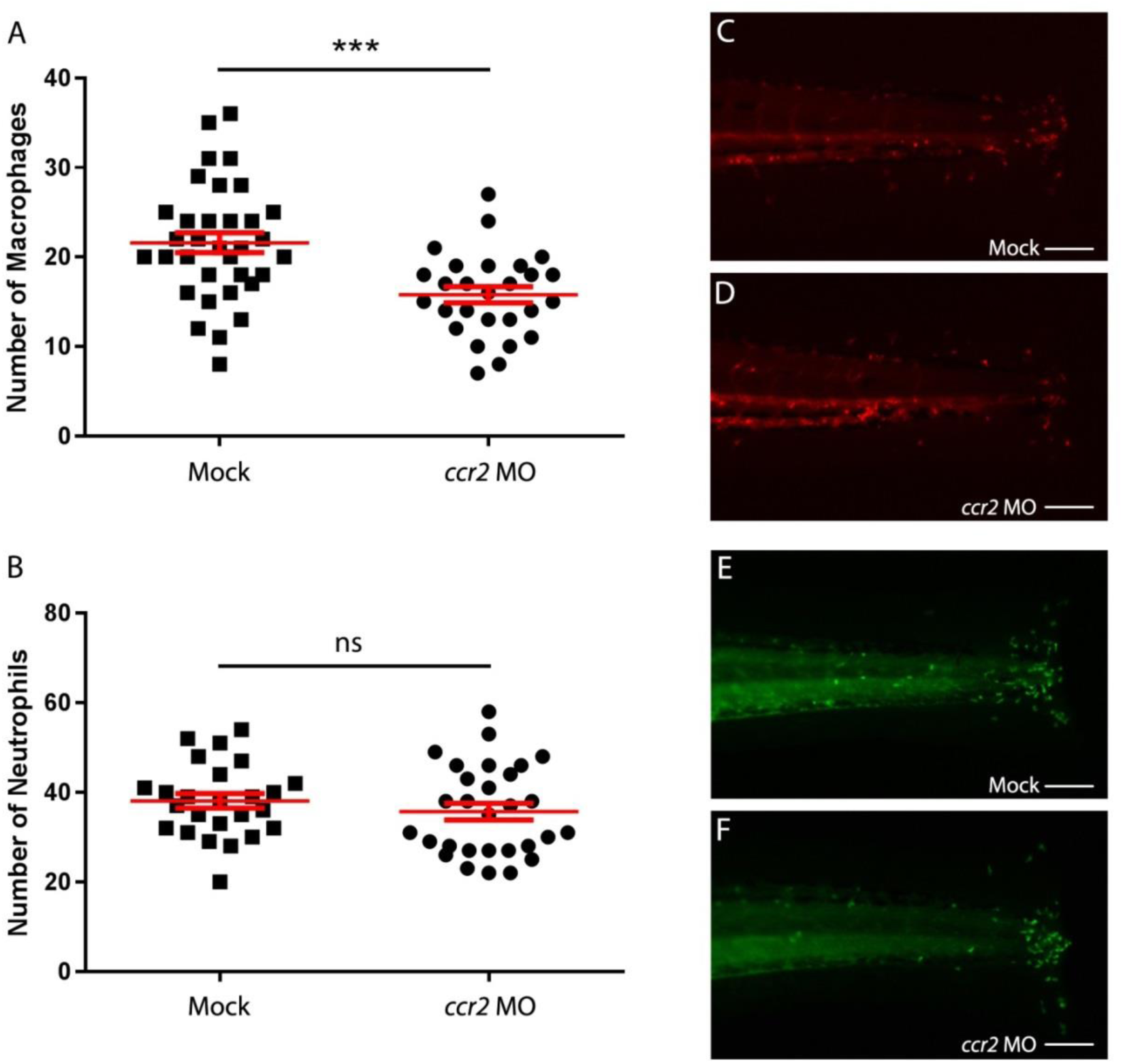
Effect of *ccr2* morpholino knockdown on macrophage and neutrophil recruitment upon tail amputation in *Tg*(*mpx:GFP/mpeg1:mCherry-F*) larvae. A-B. The number of macrophages (A) and neutrophils (B) recruited to the wounded area at 4 hpa in 3dpf larvae. In *ccr2* morphant larvae, a significantly reduced number of macrophages were recruited compared to the number in wild-type larvae. No significant difference was observed for the number of recruited neutrophils. Data were pooled from 2 independent experiments. Means ± s.e.m. are indicated in red. Statistical significance is indicated by: ns, non-significant; *** P<0.001. C-D. Representative images of the macrophages (fluorescently labeled by mCherry) (C, D) and the neutrophils (fluorescently labeled by GFP) (E, F) of wild type and *ccr2* morphant larvae at 4 hpa. Scale bar = 100 µm.

**Figure 4.**
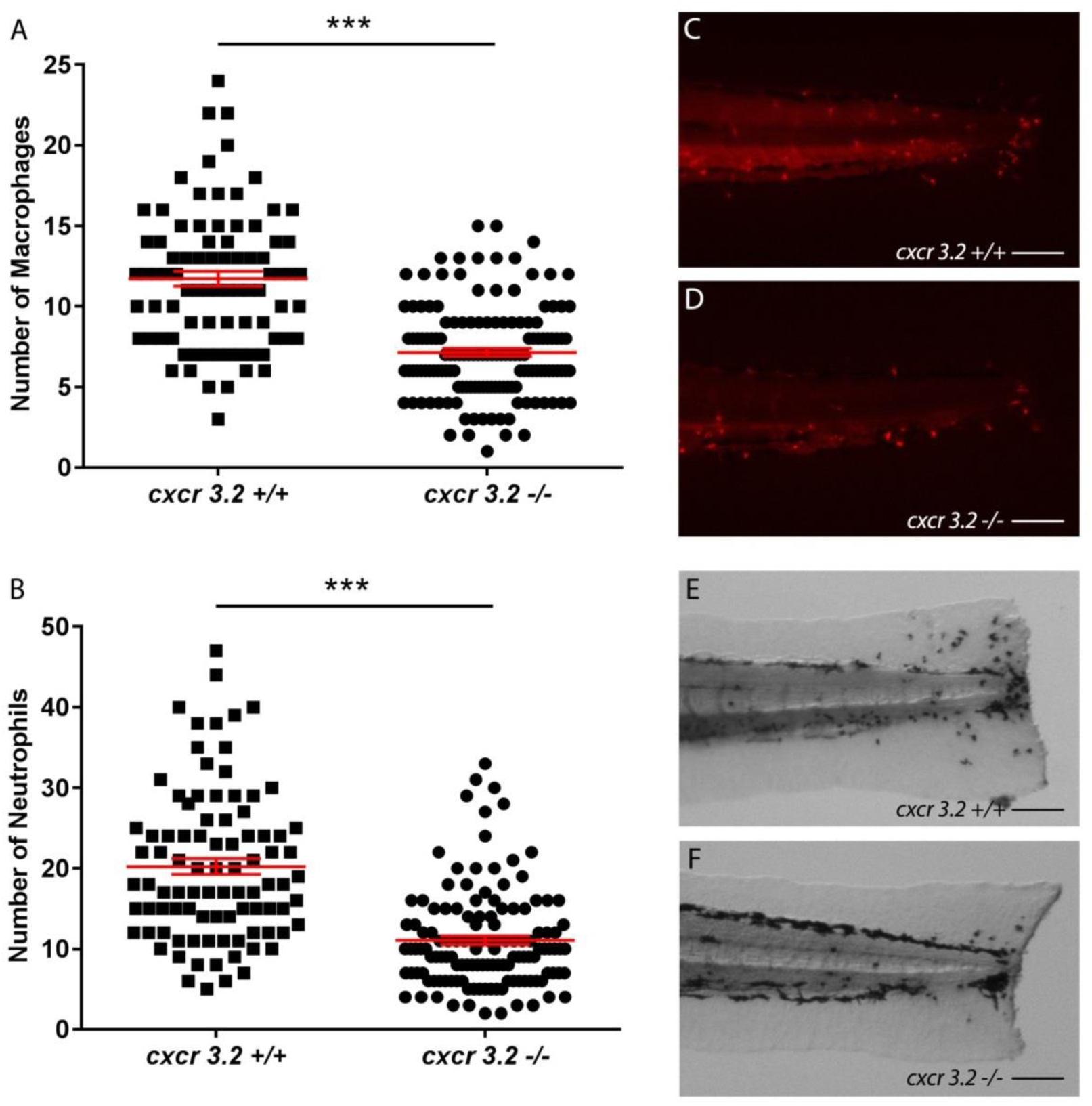
Effect of *cxcr3.2* mutation on macrophage and neutrophil recruitment upon tail amputation in *Tg*(*mpeg1:mCherry-F*) larvae. A-B. The number of macrophages (A) and neutrophils (B) that recruited to the wounded area at 4 hpa in 3 dpf larvae. A significantly reduced number of macrophages and neutrophils were recruited in *cxcr3.2* mutant larvae compared to the number in wild type controls. Data were pooled from 3 independent experiments. Means ± s.e.m. are indicated in red. Statistical significance is indicated by: *** P<0.001. C-F. Representative images of the macrophages (fluorescently labeled by mCherry) (C, D) and the neutrophils (stained using MPX assay) (E, F) of wild type and *cxcr3.2* mutant larvae at 4 hpa. Scale bar = 100 µm.

These findings indicate that beclomethasone does not affect the amputation-induced increase in the expression of the genes encoding the chemo-attractants Ccl2 and Cxcl11aa, which are specifically involved in macrophage recruitment upon tail amputation. This provides an explanation for the insensitivity of macrophage migration to glucocorticoid treatment.

### Beclomethasone attenuates almost all amputation-induced changes in gene expression in macrophages

To study whether glucocorticoid treatment changes the transcriptional response of macrophages to wounding, we performed a transcriptome analysis on FACS-sorted macrophages derived from larvae at 4 hpa. We found that 620 genes were significantly regulated by amputation, of which 411 genes were upregulated and 209 genes were downregulated (Figure 5 A, D, E). When the larvae had been amputated and treated with beclomethasone, only 327 significantly regulated genes were identified, of which 260 genes were upregulated and 67 genes were downregulated (Figure 5B, D, E).

**Figure 5.**
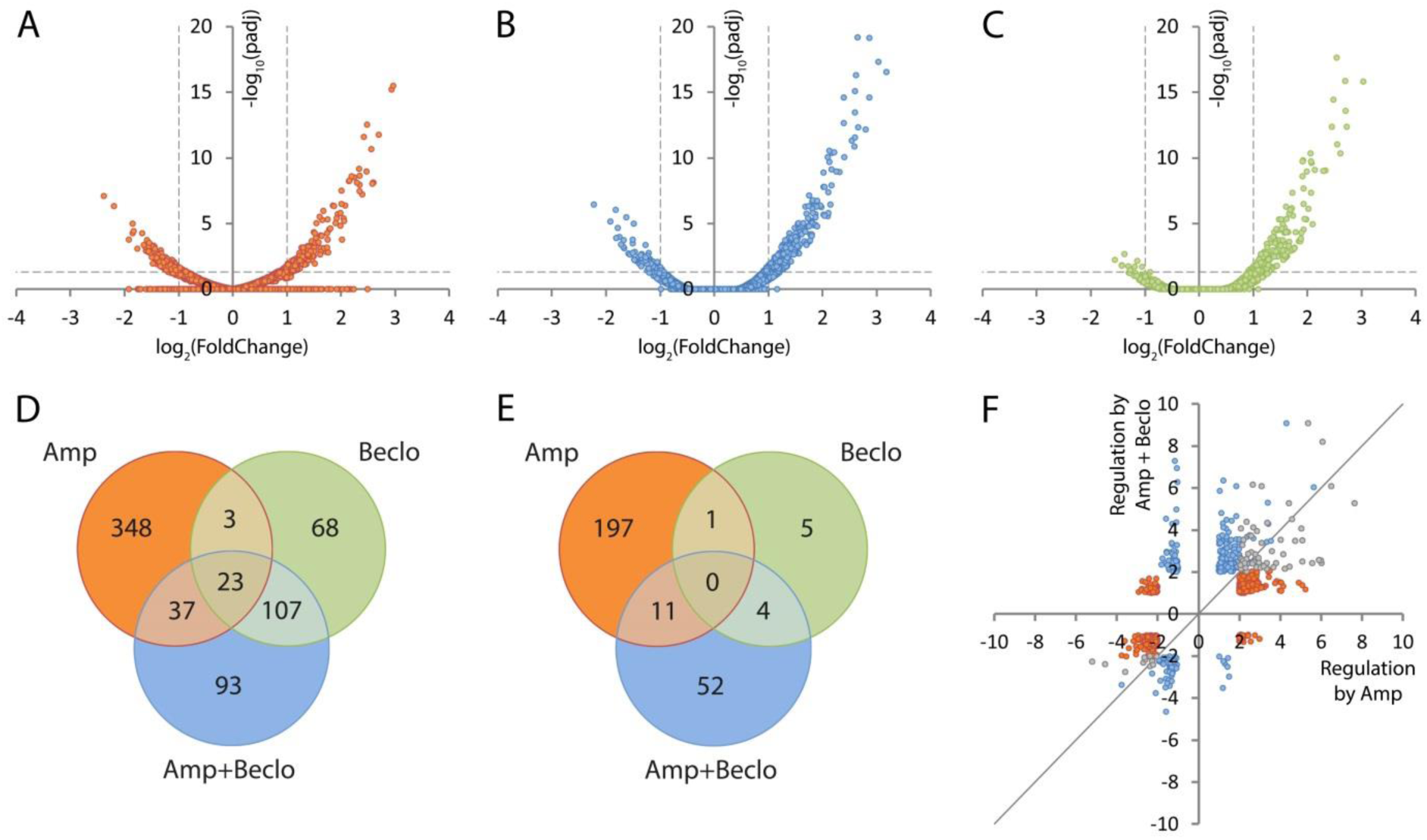
Macrophage-specific transcriptome analysis by RNA sequencing showing modulation of amputation-induced gene regulation by beclomethasone. A-C. Volcano plots indicating the fold change (x-axis) and p-value (y-axis) of the regulation for individual genes by amputation (A), beclomethasone (B) and the combined amputation/beclomethasone treatment (C). D-E. Venn diagrams showing overlaps between clusters of genes significantly upregulated (D) or downregulated (E) by amputation (Amp), beclomethasone (Beclo) and the combined amputation/beclomethasone treatment (Amp+Beclo). The diagrams show that there is a large number of genes regulated by amputation in macrophages. Beclomethasone affects the expression of a relatively small number of genes, but it decreases the number of genes significantly regulated upon amputation. F. Scatter plot showing the effect of beclomethasone treatment on amputation-regulated gene expression. For all genes showing significant regulation upon amputation (red and grey dots) or the combined beclomethasone and amputation treatment (blue and grey dots), the fold change due to beclomethasone and amputation treatment was plotted as a function of the fold change due to amputation. The grey dots represent the overlap between amputation and combination treatment. The grey line indicates the point at which beclomethasone treatment does not alter amputation-induced gene regulation. Of all the genes that were significantly regulated by amputation in macrophages, 75.37% showed attenuation in the presence of beclomethasone. Significantly regulated genes were selected by using a p.adj<0.05 and |FoldChange|>2 cutoff.

Apparently, amputation-induced gene regulation in macrophages is attenuated by beclomethasone administration. To study the effect of beclomethasone on the amputation-induced changes in gene expression in macrophages in more detail, we plotted the level of regulation by the combined amputation and beclomethasone treatment against the regulation by amputation in the absence of beclomethasone, for all genes that were significantly regulated by at least one of these treatments (Figure 5F). The resulting scatter plot shows that 75.37% of the genes regulated by amputation showed attenuation of this regulation when amputation was performed in the presence of beclomethasone. These results indicate that beclomethasone has a very general and strong dampening effect on the amputation-induced changes in gene expression in macrophages, which is in contrast with the lack of inhibition of the migration of these cells towards the wounded area.

Interestingly, only a small overlap was observed between the cluster of 620 amputation-regulated genes and the cluster of 327 genes regulated by the combined amputation and beclomethasone treatment (Figure 5 A, B, D, E). Only 60 and 11 genes were present in the overlap between these clusters for up-regulation and down-regulation respectively (Figure 5 D, E). A large overlap was observed between the gene cluster regulated by the combination treatment and the cluster regulated by beclomethasone (without amputation) (134 genes in total, Figure 5D, E, Supplementary Figure 1A). This indicates that the cluster of genes regulated by the combination treatment mainly contains genes that are regulated as a result of the beclomethasone treatment. Apparently, amputation hardly affects beclomethasone-induced changes in gene expression, whereas beclomethasone has a very strong effect on amputation-induced transcriptional changes.

Using gene ontology analysis, we classified the regulated genes according to the KEGG-pathways they are involved in (Supplementary Figure 2, Supplementary Table 1). This analysis showed that the largest group of pathways regulated by amputation were involved in metabolism (16 pathways, 98 genes) and that 4 pathways (19 genes) involved in the immune system were altered. The combined amputation and beclomethasone treatment affected a smaller number of pathways for both metabolism- and immune system-related pathways (12 pathways and 26 genes, and 1 pathway and 6 genes respectively). Only 5 of these pathways (Toll-like receptor signaling pathway, Insulin resistance, Biosynthesis of antibiotics, Galactose metabolism, Glycolysis/Gluconeogenesis) were both regulated by amputation and by the combination treatment. Beclomethasone treatment (without amputation) affected 7 pathways (5 metabolism-related), of which 6 were also regulated when the larvae were amputated in the presence of beclomethasone.

Among the significantly enriched metabolism-related KEGG pathways, we studied 3 specific pathways which are known to be associated with specific macrophage phenotypes: glycolytic metabolism which is increased in pro-inflammatory (M1) macrophages, and mitochondrial oxidative phosphorylation (OXPHOS) and tricarboxylic acid (TCA) cycle which are related to the anti-inflammatory (M2) phenotype (Van den Bossche et al., 2015). We mapped the gene expression levels into these pathways (Supplementary Figure 3 A-C). The data showed that the vast majority of the mapped genes were up-regulated by amputation and this up-regulation was inhibited by beclomethasone treatment. We, therefore, conclude from the gene ontology analysis that amputation mainly up-regulates genes involved in metabolism and the immune system and that the vast majority of the amputation-induced changes in these gene ontology groups are attenuated by glucocorticoids.

### Glucocorticoids inhibit the differentiation of macrophages towards a pro-inflammatory phenotype

Subsequently, we specifically analyzed the regulation of immune-related genes. For all immune-related genes that were significantly regulated by amputation, we plotted the regulation by amputation, by beclomethasone, and by the combination of amputation and beclomethasone (Figure 6). For the vast majority of these genes, the amputation-induced changes were attenuated by the administration of beclomethasone. Among those genes were 3 that are known to be associated with a pro-inflammatory (M1) phenotype of macrophages: *tnfa*, *il1b*, and *il6* (Martinez and Gordon, 2014; Nguyen-Chi et al., 2015). For 3 genes (*cd22*, *alox5ap*, and *tlr5b*), the amputation-induced regulation was enhanced by beclomethasone. These findings suggest that the differentiation of macrophages to a pro-inflammatory (M1) phenotype is sensitive to inhibition by glucocorticoids.

**Figure 6.**
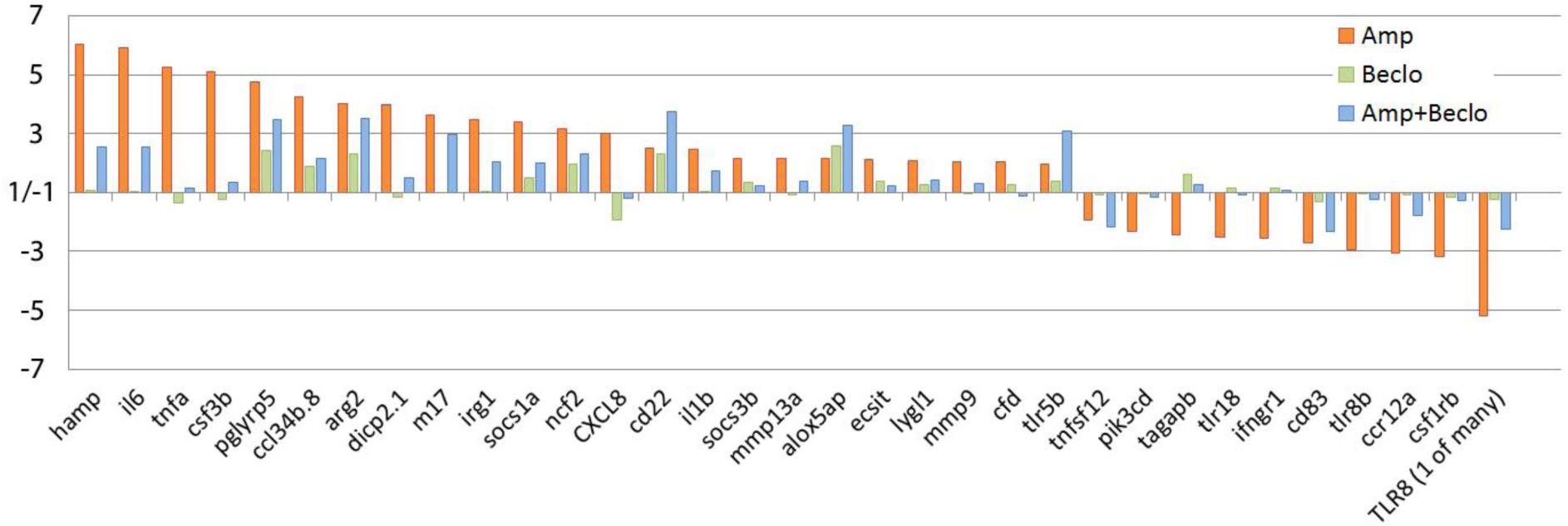
Regulation of immune-related genes in macrophages, determined by RNA sequencing analysis. For all genes significantly regulated upon amputation, the fold change due to amputation (Amp; red bars), beclomethasone (Beclo; green bars), and the combined amputation/beclomethasone treatment (Amp+Beclo; blue bars) is shown. The results show that beclomethasone dampens the amputation-induced expression of most genes, but for 4 genes (*cd22, alox5ap, tlr5b*) the combined treatment results in a higher fold change compared to the amputation treatment.

To study the glucocorticoid sensitivity of macrophage differentiation in more detail and validate some of the observed transcriptional changes, we performed qPCR on RNA samples isolated from FACS-sorted macrophages. At 4 hpa, the expression of 4 classic pro-inflammatory genes was measured: *il6, il1b, tnfa*, *mmp9*, of which the first 3 are markers for M1 macrophages and the 4^th^ encodes a metalloproteinase that facilitates leukocyte migration by remodeling the extracellular matrix (Martinez and Gordon, 2014; Nguyen-Chi et al., 2015; Rohani and Parks, 2015) (Figure 7A). The expression levels of *il6* and *il1b* showed an amputation-induced increase, and this increase was attenuated upon the combined beclomethasone and amputation treatment. The levels of *tnfa* and *mmp9* expression were not significantly increased by amputation, but the expression level of *tnfa* was significantly lower after the combination treatment compared to the amputation treatment.

**Figure 7.**
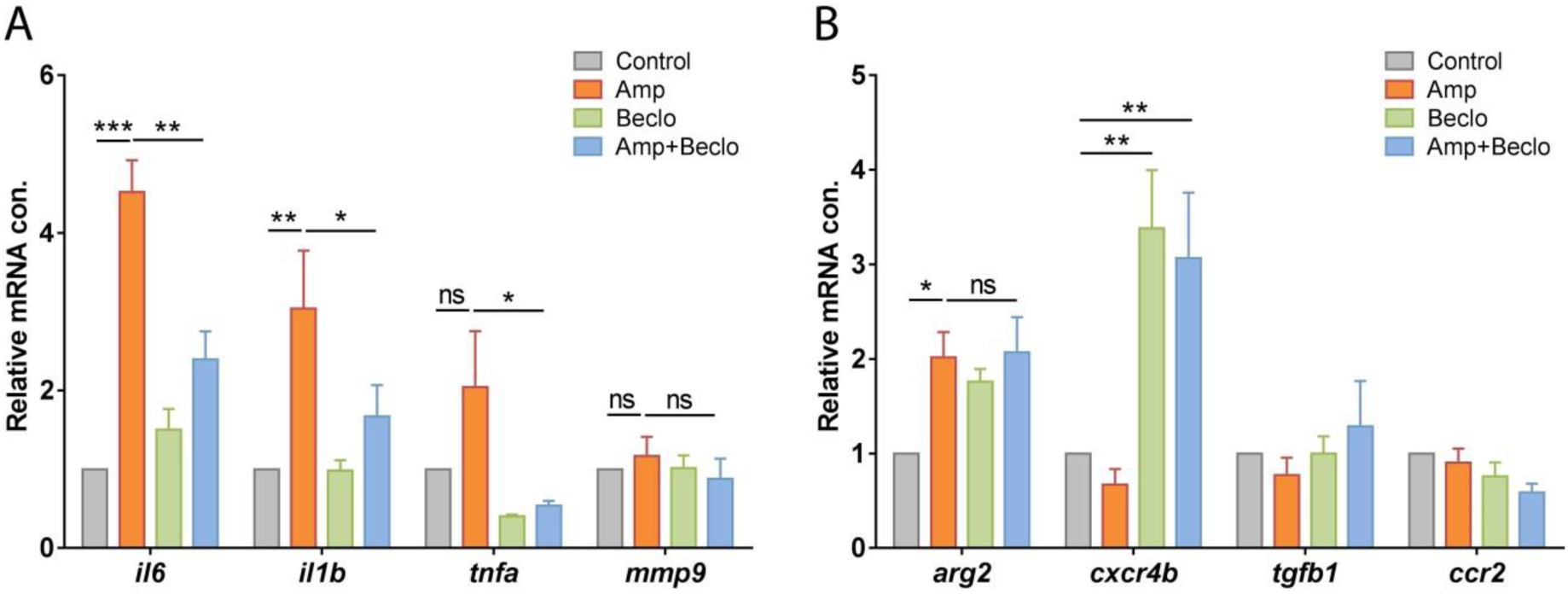
Expression levels of immune-related genes in FACS-sorted macrophages, determined by qPCR for *il6*, *il1b*, *tnfa, mmp9* (A) and for *arg2, cxcr4b, tgfb1, ccr2* (B) at 4hpa in 3 dpf larvae. The levels of *il6* and *il1b* expression were significantly increased by amputation, and this effect was inhibited by beclomethasone treatment. The expression level of *arg2* showed a significant increase upon amputation, and beclomethasone treatment did not affect this regulation. The expression level of *cxcr4b* was increased by beclomethasone treatment. Values shown are the means ± s.e.m. of three independent experiments. Statistical significance is indicated by: ns, non-significant; * P<0.05; ** P<0.01; *** P<0.001.

In addition, we measured the expression levels of 4 markers for M2 macrophages, *arg2, cxcr4b, tgfb1* and *ccr2* (Nguyen-Chi et al., 2015; Yang and Ming, 2014)(Figure 7B). The expression level of *arg2* was induced by amputation at 4 hpa, and this induction was similar upon the combination treatment. The other genes were not upregulated by amputation at this time point, but upon beclomethasone treatment the expression of *cxcr4b* was increased. Since the M2 macrophage markers are expected to show increased expression levels during the resolution phase of the inflammatory response (Nguyen-Chi et al., 2015), we measured the expression of those genes in macrophages at 24 hpa as well (Supplementary Figure 4A). However, no significant upregulation by amputation was observed for any of these 4 genes. Thus, in this experiment on M2 markers, we only found an amputation-induced upregulation of the expression of *arg2* at 4 hpa, and this upregulation was insensitive to beclomethasone.

To further study the influence of beclomethasone on the differentiation of macrophages towards a pro-inflammatory (M1) phenotype, we used a reporter line for the expression of *tnfa*: the *Tg(mpeg1:mCherry-F/tnfa:eGFP-F)* fish line. Larvae from this line were amputated at 5 dpf, and at a more distal position than in the previous experiments to create a wound that recruits fewer macrophages which facilitates the visualization of individual *tnfa* expressing macrophages. We performed live confocal imaging at 2 and 4 hpa, and the GFP expression level in macrophages was used as a reporter for *tnfa* promoter activity *in vivo* (Figure 8 A-C). In the control group, an increase in the percentage of GFP-positive macrophages was observed between 2 and 4 hpa. During this time course, this percentage increased from 9.8±3.4% to 23.8±4.0%. In the beclomethasone-treated group, at both time points, a lower percentage of *tnfa* expressing macrophages was recruited to the wounded area compared to the control group (1.7±1.7% and 1.4±1.4% respectively).

**Figure 8.**
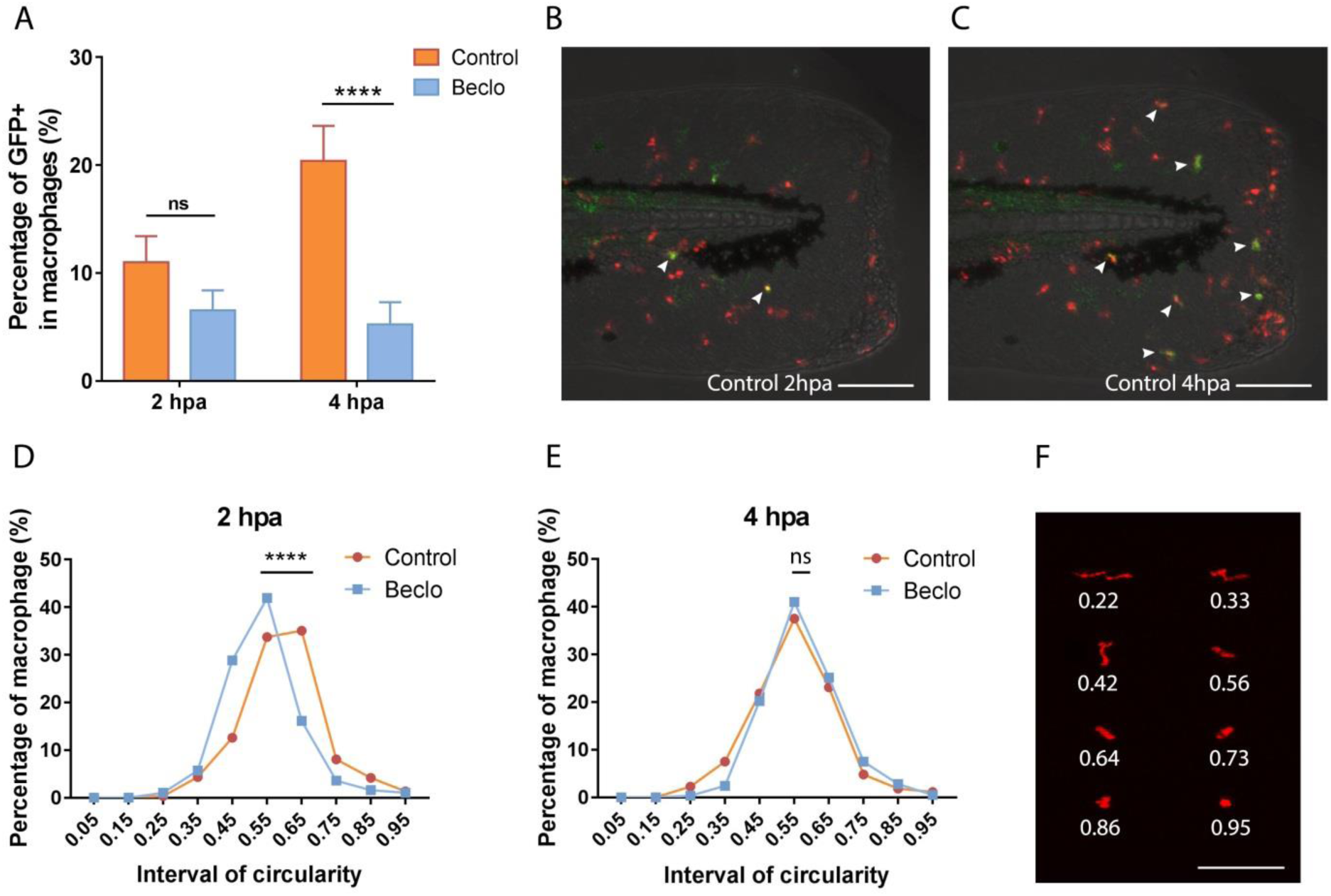
Effect of beclomethasone on the differentiation of macrophages. GFP expression in *Tg(mpeg1:mCherry-F/tnfa:eGFP-F)* reporter larvae and less elongated and dendritic morphology are considered markers for differentiation towards a pro-inflammatory (M1) phenotype. A. The number of GFP-positive macrophages (as a percentage of the total number of macrophages) recruited to the wounded area were quantified at 2 and 4 hpa in 5 dpf larvae. In the beclomethasone-treated group, at 4 hpa, a significantly reduced percentage of the recruited macrophages was GFP-positive compared to the control group. Values shown are means ± s.e.m. Statistical significance is indicated by: ns, non-significant; **** P<0.0001. B-C. Representative images of macrophages (fluorescently labeled by mCherry) and GFP-positive macrophages (fluorescently labeled by both mCherry and GFP) in the control group at 2 hpa (B) and 4 hpa (C). Scale bar = 100 µm. Arrow heads indicate macrophages displaying the GFP signal, which is a measure for activation of the *tnfa* promoter. D-E. The distribution of circularity of macrophages recruited to the wounded area at 2 hpa (D) and 4 hpa (E) in 3 dpf larvae. At 2 hpa, a significant difference of distribution pattern was observed between the two groups, with the beclomethasone-treated group shifted towards lower circularity. At 4 hpa, no significant difference was observed. Statistical significance is indicated by: ns, non-significant; **** P<0.0001. F. Representative images of macrophages analyzed in D and E and their corresponding circularity. Scale bar = 100 µm.

Finally, we analyzed the influence of beclomethasone on the morphology of macrophages, since macrophage morphology has been shown to be an indicator for their differentiation: M1 macrophages are less elongated and dendritic than M2 macrophages (Nguyen-Chi et al., 2015). A small hole was punched on the tail fins of the larvae with a glass microcapillary needle (Benard et al., 2012) to recruit a small number of leukocytes for visualization of individual cells. We performed live confocal imaging at 3 dpf with the *Tg(mpx:GFP/mpeg1:mCherry-F)* fish line and the circularity of mCherry-positive macrophages was used to quantitate the morphology (Figure 8 D-E). In the control group, at 2 hpa, the percentage of macrophages with a high circularity (0.5-1.0) was relatively high (67.6±4.0%) and gradually decreased to 47.9±3.2% at 12 hpa (Supplementary Figure 5 A). In the beclomethasone-treated group, at 2 hpa the percentage of macrophages with a high circularity was lower (51.7±3.5%) and remained relatively stable until 12 hpa (Supplementary Figure 5 B). The most obvious difference between the control and beclomethasone-treated group was observed at 2 hpa. At this time point, the plot showing the distribution of circularity shows a clear shift towards a lower circularity in the beclomethasone-treated group (Figure 8 D). At 4 hpa, this difference of circularity distribution between the control group and beclomethasone-treated group had disappeared (Figure 8 E). Thus, both the data obtained using the *tnfa:eGFP-F* reporter line and the data from the analysis of the circularity showed an inhibitory effect of beclomethasone on the differentiation of macrophages towards a pro-inflammatory (M1) phenotype.

## Discussion

Although glucocorticoids have been used as anti-inflammatory drugs for decades, their mechanism of action and the specificity of their effects have not been fully unraveled yet. Using the zebrafish tail amputation model, we have shown that the inflammatory response comprises glucocorticoid-sensitive and glucocorticoid-insensitive pathways. Glucocorticoids inhibit the migration of neutrophils towards a site of inflammation by inhibiting the induction of chemoattractants for this cell type. However, the migration of macrophages is not affected by glucocorticoids, since the induction of two chemoattractants that are critical for macrophage recruitment, *ccl2* and *cxcl11aa*, is insensitive to glucocorticoid treatment. Using RNAseq analysis we show that the glucocorticoid beclomethasone attenuates most transcriptional responses to amputation in macrophages and inhibit their differentiation towards a pro-inflammatory (M1) phenotype.

Chemoattractants are important trafficking signals that direct the movement of immune cells into and out of specific tissues (Luster et al., 2005). In this study, we have demonstrated that glucocorticoids exert a specific inhibitory effect on the induction of the expression of two chemoattractants involved in neutrophil recruitment (Il8 and Cxcl18b). Using *in vitro* and *in vivo* models, it has been demonstrated that human and mouse neutrophil migration is dependent on the induction of Il8 expression (Godaly et al., 2000; Huber et al., 1991; Kaunisto et al., 2015) and that this induction is inhibited by glucocorticoids (Huang et al., 2015; Keelan et al., 1997; Yano et al., 2006). In mammals, Il8 has been demonstrated to signal through the chemokine receptors Cxcr1 and Cxcr2, whereas in zebrafish only Cxcr2 has been shown to mediate the effects of Il8 (Brugman, 2016; Torraca et al., 2015). Cxcl18b, a chemokine specific for fish and amphibian species, has also been shown to act as a ligand for Cxcr2 in zebrafish, thereby stimulating chemotaxis of neutrophils (Torraca et al., 2017). These findings suggest that Cxcr2 activation is crucial for the migration of neutrophils and that glucocorticoids inhibit this migration by attenuating the induction of the expression of Cxcr2 agonists like Il8 and Cxcl18b.

In contrast to the inhibitory effect on neutrophil migration, our study revealed that glucocorticoids leave the induction of chemoattractants involved in macrophage recruitment (Ccl2 and Cxcl11aa) unaffected. Ccl2, also known as monocyte chemoattractant protein-1 (MCP-1), and Cxcl11aa have been shown to be key chemokines implicated in macrophage migration and infiltration in humans and mice (Deshmane et al., 2009; Gunn et al., 1997; Lee et al., 2018; Shen et al., 2014; Szebeni et al., 2017; Wada et al., 1999). In zebrafish, their role as chemoattractants for macrophages has been demonstrated during mycobacterial infection (Cambier et al., 2017; Cambier et al., 2014; Torraca et al., 2015). Our data show that these two chemoattractants also promote macrophage migration in the tail amputation model and that beclomethasone has no effect on the amputation-induced increase in their expression levels. These results may provide a mechanistic explanation for the glucocorticoid insensitivity of macrophage-dominated inflammatory diseases like COPD. In line with our findings, it has been shown in bronchoalveolar lavage fluid of COPD patients that glucocorticoid treatment reduces neutrophil numbers, but that the number of macrophages was not decreased (Jen et al., 2012).

Contrary to our findings, in most of the studies carried out in humans and rats, the inflammation-induced Ccl2 level has been found to be inhibited by glucocorticoids (Kim et al., 1995; Little et al., 2006; Wada et al., 1999), and this inhibition is related to a decreased p38 MAPK phosphorylation (Baldassare et al., 1999; Little et al., 2006). Similarly, glucocorticoids have been shown to inhibit Cxcl11 upregulation in fluticasone propionate-stimulated peripheral blood monocytes, and in INF-γ- or LPS-stimulated RAW 264.7 macrophages, as well as in multiple tissues of endotoxemia mice (Ehrchen et al., 2007; Widney et al., 2000). Nevertheless, some studies do show an insensitivity of the mammalian Ccl2 or Cxcl11aa induction to glucocorticoid treatment. In a breast cancer cell line (T47D), glucocorticoid treatment has no effect on Il1-stimulated Ccl2 production (Kelly et al., 1997), and in A579 epithelial cells, IFNγ-induced Cxcl11 is insensitive to glucocorticoid treatment (O’Connell et al., 2015b). These data suggest that the observed insensitivity of the *ccl2* and *cxcl11a* induction to glucocorticoids, which underlies the glucocorticoid insensitivity of macrophage migration, requires a specific context, which may involve factors like the activating signal, the glucocorticoid treatment regime, or the cell type and tissue involved.

Although glucocorticoids did not affect the migration of macrophages in our study, they did have a big impact on the transcriptional changes in these cells upon amputation. We showed by RNAseq analysis in FACS-sorted macrophages that, similarly to our previous findings from a microarray analysis carried out on RNA isolated from whole larvae (Chatzopoulou et al., 2016), most of the amputation-induced transcriptional changes are decreased by beclomethasone, whereas a small subset of transcriptional responses is insensitive to glucocorticoid treatment. Focusing on the regulation of immune-related genes, we found that, in line with our previous findings in whole larvae (Chatzopoulou et al., 2016), beclomethasone suppressed the induction of almost all pro-inflammatory genes, like *il6*, *tnfa*, *il1b*, *il8* and *mmp9*. In line with these data, many genes involved in glycolysis, a metabolic pathway often associated with an M1 phenotype (Kelly and O’Neill, 2015; Saha et al., 2017; Van den Bossche et al., 2015), were upregulated upon amputation and this upregulation was mostly inhibited by beclomethasone. This inhibitory effect of glucocorticoids on the induction of M1-associated genes in macrophages is in agreement with *in vitro* results obtained in LPS-stimulated primary mouse macrophages (Ogawa et al., 2005; Sacta et al., 2018; Uhlenhaut et al., 2013), and *in vivo* data obtained in a mouse model for arthritis (Hofkens et al., 2013). In addition, by *in vivo* imaging of the wound-induced inflammatory response, we observed a reduction in the number of macrophages with activation of a *tnfa:eGFP-F* reporter gene and a morphology characterized by a low circularity, which demonstrates that the macrophage differentiation to an M1 phenotype was inhibited. Taken together, these data strongly support the idea that glucocorticoids inhibit the differentiation of macrophages to an M1 phenotype by interfering at the level of transcription.

In addition to the effect of glucocorticoids on M1 differentiation, we investigated their effect on the differentiation of macrophages to an M2 phenotype. Previous studies in a mouse arthritis model showed that the induction of an M2 phenotype was not affected by glucocorticoids (Hofkens et al., 2013) and in an acute lung injury model (Tu et al., 2017) it was shown to be enhanced. In our RNAseq and qPCR analysis, the typical M2 marker *arg2* (Martinez and Gordon, 2014; Nguyen-Chi et al., 2015; Yang and Ming, 2014) was among the small subset of amputation-induced genes that were insensitive to beclomethasone, suggesting that the differentiation to an M2 phenotype is insensitive to glucocorticoids. However, genes involved in the TCA cycle and OXPHOS, metabolic pathways associated with an M2 phenotype (Kelly and O’Neill, 2015; Saha et al., 2017; Van den Bossche et al., 2015), were upregulated upon amputation and this upregulation was inhibited by beclomethasone, which would suggest that M2 differentiation is blocked by glucocorticoid treatment. In our qPCR analysis, we showed that the expression of various M2 markers (*cxcr4b, tgfb1, ccr2*) was not increased upon amputation, and this may indicate that macrophages do not fully develop the M2 phenotype in our assay. In summary, whereas the amputation-induced increases in the expression levels of M1 markers are consistently inhibited by glucocorticoids, increased expression of M2 markers (when present in our assay) can be either insensitive or suppressed.

In our tail amputation model for inflammation, the vast majority of transcriptional responses was suppressed by glucocorticoids and only a small subset of these responses was not affected. These data may indicate an important (but not exclusive) role for the transcription factor NF-κB in the observed inflammatory response, since in many studies it has been shown that the NF-κB-mediated transcriptional activation can be suppressed by glucocorticoids or remain unaffected (Kuznetsova et al., 2015; Ogawa et al., 2005; Rao et al., 2011; Sacta et al., 2018). Recruitment of IRF3 to the transcription initiation complex has been shown to be associated with sensitivity to GR suppression (Ogawa et al., 2005; Uhlenhaut et al., 2013). The activities of some other transcription factors are differently modulated upon glucocorticoid treatment, which suggests that they play a minor role in the inflammatory response in our assay. For example, the IFNγ-induced STAT1 activity and the poly(dA:dT)-induced activation of the cytosolic DNA sensing pathway have been shown to be insensitive to glucocorticoids (O’Connell et al., 2015a; Wang et al., 2017), and the LIF/IL-6-induced STAT3 activity has even been found to be enhanced by glucocorticoids (Anna et al., 2012; Busillo and Cidlowski, 2013; Langlais et al., 2008; Langlais et al., 2012). The latter pathway may play a role in the enhancement of the inflammatory response by glucocorticoids observed in zebrafish larvae with a deficiency in the hematopoietic phosphatase Ptpn6/Shp1 (Kanwal et al., 2013).

In conclusion, our *in vivo* study of the glucocorticoid modulation of the transcriptional response to wounding using the zebrafish tail amputation model shows that the vast majority of these responses are sensitive to glucocorticoids and a small subset is insensitive. These insensitive responses are involved in the migration of macrophages and may provide an explanation for the glucocorticoid resistance observed in macrophage-dominated inflammatory diseases like COPD. The sensitive responses are involved in the differentiation of macrophages to an M1 phenotype and the migration of neutrophils. The characterization of these pathways and increased understanding of the molecular mechanisms underlying the differences in glucocorticoid sensitivity will be crucial for the future development of novel anti-inflammatory glucocorticoid therapies.

## Materials and methods

### Zebrafish lines and maintenance

Zebrafish were maintained and handled according to the guidelines from the Zebrafish Model Organism Database (http://zfin.org) and in compliance with the directives of the local animal welfare committee of Leiden University. They were exposed to a 14 hours light and 10 hours dark cycle to maintain circadian rhythmicity. Fertilization was performed by natural spawning at the beginning of the light period. Eggs were collected and raised at 28°C in egg water (60 µg/ml Instant Ocean sea salts and 0.0025% methylene blue).

The following fish lines were used in this work: wild type (wt) strain AB/TL, the double transgenic lines *Tg(mpx:GFP/mpeg1:mCherry-F)* (Bernut et al., 2014; Renshaw et al., 2006) and *Tg(mpeg1:mCherry-F/tnfa:eGFP-F)* (Nguyen-Chi et al., 2015), and the combination of *Tg(mpeg1:mCherry-F)* and the homozygous mutants (*cxcr3.2*^−/−^) or wt siblings (*cxcr3.2*^+/+^) of the *cxcr3.2^hu6044^* mutant strain(Torraca et al., 2015).

### Tail amputation and chemical treatments

After anesthesia with 0.02% aminobenzoic acid ethyl ester (tricaine, Sigma Aldrich), the tails of 3 days post fertilization (dpf) embryos were partially amputated (Figure 1A) with a 1mm sapphire blade (World Precision Instruments) on 2% agarose-coated Petri dishes under a Leica M165C stereomicroscope (Chatzopoulou et al., 2016). Amputated and non-amputated (control) embryos were pretreated for 2 hours with 25 µM beclomethasone (Sigma Aldrich) or vehicle (0.05% dimethyl sulfoxide (DMSO)) in egg water prior to amputation and received the same treatment after the amputation.

### Confocal microscopy and image analysis

The amputated larvae were mounted in 1.2% low melting agarose in egg water containing 0.02% tricaine and 25 µM beclomethasone or 0.05% DMSO on 40 mm glass bottom dishes (Willco-Dish) and covered with 1.5 ml egg water containing tricaine and the corresponding chemicals. Confocal microscopy was performed using a Nikon Eclipse Ti-E microscope with a Plan Apo 20X/0.75 NA objective. A 488 nm laser was used for excitation of GFP and a 561 nm laser was used for excitation of mCherry. Time-lapse microscopy was performed at 28 °C with an interval of approximately one minute. From the obtained z-stacks, aligned maximum projection images were generated using NIS-Elements, which were further analyzed using Image J with plugins “Local Maxima Stack” and “Track Foci” developed by Dr. Joost Willemse (Leiden University).

### Morpholino injection

A morpholino targeting the translational start site of the *ccr2* gene (5’AACTACTGTTTTGTGTCGCCGAC3’, purchased from Gene Tools) (Cambier et al., 2014) was prepared and stored according to the manufacturer’s instructions. Injection of 1 nl (0.5 mM) of the morpholino solution was performed into the yolk of fertilized eggs at the 1-2 cell stage.

### RNA isolation, cDNA synthesis and quantitative PCR (qPCR)

At different time points after amputation, larvae were collected (15-20 per sample) in QIAzol lysis reagent (QIAGEN) for RNA isolation, which was performed using the miRNeasy mini kit (Qiagen), according to the manufacturer’s instructions. Extracted total RNA was reverse-transcribed using the iScript™ cDNA Synthesis Kit (Bio-Rad). QPCR was performed on a MyiQ Single-Color Real-Time PCR Detection System (Bio-Rad) using iTaq™ Universal SYBR^®^ Green Supermix (Bio-Rad). The sequences of the primers used are provided in Supplementary Table 2. Cycling conditions were pre-denaturation for 3 min at 95°C, followed by 40 cycles of denaturation for 15 s at 95°C, annealing for 30 s at 60°C, and elongation for 30 s at 72°C. Fluorescent signals were detected at the end of each cycle. Cycle threshold values (Ct values, i.e. the cycle numbers at which a threshold value of the fluorescence intensity was reached) were determined for each sample. For each sample, the Ct value was subtracted from the Ct value of a control sample, and the fold change of gene expression was calculated and adjusted to the expression levels of a reference gene (*peptidylprolyl isomerase Ab* (*ppiab*)). Data shown are means ± s.e.m. of three independent experiments.

### Myeloperoxidase (Mpx) staining

Larvae were fixed in 4% paraformaldehyde (PFA, Sigma Aldrich) at 4°C overnight, and rinsed with PBS containing 0.05% Tween20. The myeloperoxidase (mpx) staining for the *cxcr3.2* mutant line was performed using the Peroxidase (Myeloperoxidase) Leukocyte kit (Sigma Aldrich), according to the manufacturer’s instructions. To visualize both macrophages and neutrophils in the same larvae, the mpx staining was always performed after imaging of the fluorescent signal of the macrophages.

### Imaging and image quantification

Images of fixed or live larvae were captured using a Leica M205FA fluorescence stereomicroscope, equipped with a Leica DFC 345FX camera. In all fish lines used, the macrophages were detected based on the fluorescence of their mCherry label. Neutrophils were detected based on either their fluorescent GFP label or their mpx staining. To quantify the number of macrophages and/or neutrophils recruited to the wounded area, the cells in a defined area of the tail (Figure 1A) were counted manually. Data were pooled from two or three independent experiments, and the means ± s.e.m. of the pooled data are indicated.

### Fluorescence-Activated Cell Sorting (FACS) of macrophages

Macrophages were sorted from *Tg(mpeg1.4:mCherry-F)* embryos as previously described (Rougeot et al., 2014; Zakrzewska et al., 2010). Dissociation was performed with 100-150 embryos for each sample at 4 hours post amputation (hpa) using Liberase TL (Roche) and stopped by adding Fetal Calf Serum (FCS) to a final concentration of 10%. Isolated cells were resuspended in Dulbecco’s PBS (DPBS), and filtered through a 40 µm cell strainer. Actinomycin D (Sigma Aldrich) was added (final concentration of 1 µg/ml) to each step to inhibit transcription. Macrophages were sorted based on their red fluorescent signal using a FACSAria III cell sorter (BD Biosciences). The sorted cells were collected in QIAzol lysis reagent (Qiagen) for RNA isolation. Extracted total RNA was either reverse-transcribed for qPCR or amplified using the SMART-seq V4 kit (Clontech) for sequencing.

### Transcriptome analysis

A total of 12 samples (four experimental groups obtained from three replicate experiments) were processed for transcriptome analysis using cDNA sequencing. The RNA seq libraries generated with the SMART-seq V4 kit were sequenced using an Illumina HiSeq 2500 instrument according to the manufacturer’s instructions with a read length of 50 nucleotides. Image analysis and base calling were done by the Illumina HCS version 2.2.68, and RTA version 1.18.66. cDNA sequencing data were analyzed by mapping the reads to the Danio rerio GRCz10 reference genome with annotation version 80 using Tophat (v2.1.0). Subsequently, the DESeq (v1.26.0) R package was used to test for differential expression. Before each analysis, the genes with low reads were removed (i.e. those genes for which the sum of reads from three replicates of the analyzed two groups was lower than 30). The output data were used for transcriptome analysis. Significant gene regulation was defined by using p.adj<0.05 and |FoldChange|>2 cutoffs. The raw data are available at Gene Expression Omnibus database under accession number GSE122643.

Gene ontology analysis was performed using the online functional classification tool Database (DAVID; http://david.abcc.ncifcrf.gov/summary.jsp) for Annotation, Visualization, and Integrated Discovery. Further analysis of the macrophage transcriptomes was performed in R v3.4.3 using *Bioconductor* v3.6. Zebrafish Ensembl gene IDs were converted to Entrez Gene IDs using the R package *org.Dr.eg.db* v3.5.0. The enriched pathways in different groups were determined by comparing the statistically differentially expressed genes against the KEGG zebrafish database using the *kegga*() function from the *edgeR* package v3.20.7. Finally, gene expression data were mapped into significantly enriched KEGG pathways using *pathview* v1.18.0.

### Statistical analysis

Statistical analysis was performed using GraphPad Prism 7 by ANOVA with a Fisher’s LSD post hoc test (Figure 1B-E, Figure2, Figure 7, Figure 8 A, Supplementary Figure 4), Kolmogorov-Smirnov test (Figure 1F, G, Figure 8 D, E) or two-tailed t-test (Figure 3, 4). Significance was accepted at p<0.05 and different significance levels are indicated: * p<0.05; ** p<0.01; *** p<0.001; **** p<0.0001.

## Acknowledgements

We thank Dr. Tomasz Prajsnar and Dr. Gabriel Forn-Cuní for their help during the transcriptomic data analysis.

## Competing interests

The authors declare no competing interests.

## Funding

Yufei Xie was funded by a grant from the China Scholarship Council (CSC).

